# Alzheimer’s Genetic Risk Score linked to Incident Mild Behavioral Impairment

**DOI:** 10.1101/200840

**Authors:** Shea J. Andrews, Zahinoor Ismail, Kaarin J. Anstey, Moyra Mortby

**Affiliations:** Centre for Research on Ageing, Health and Wellbeing, Australian National University, Canberra, Australia.; Departments of Psychiatry and Clinical Neurosciences, Mathison Centre for Mental Health Research & Education, Ron and Rene Centre for Healthy Brain Aging Research, Hotchkiss Brain Institute, University of Calgary, Calgary, Canada

## Abstract

Mild Behavioral Impairment (MBI) describes the emergence of later-life Neuropsychiatric Symptoms (NPS) as an at-risk state for cognitive decline and dementia and as a potential manifestation of prodromal dementia. How NPS mechanistically link to the development of Mild Cognitive Impairment (MCI) and Alzheimer’s disease (AD) is not fully understood. Potential mechanisms include either shared risk factors that are related to both NPS and cognitive impairment, or AD pathology promoting NPS. This is the first study to examine whether AD genetic loci, individually and as a genetic risk score, are a shared risk factor with MBI. 1377 older adults (aged 72-79; 738 males; 763 normal cognition) from the PATH Through Life project. MBI was assessed in accordance with Criterion 1 of the ISTAART-AA diagnostic criteria using the Neuropsychiatric Inventory. 25 LOAD risk loci were genotyped and a weighted genetic risk score (GRS) was constructed. Binomial logistic regression adjusting for age, gender, and education examined the association between LOAD GRS and MBI domains. An increase in the LOAD GRS and APOE*ε4 were associated with higher likelihood of Affective Dysregulation; *MS4A4A*-rs4938933*C and *MS4A6A*-rs610932*G were associated with a reduced likelihood of Affective Dysregulation; *ZCWPW1*-rs1476679*C was associated with a reduced likelihood of Social Inappropriateness and Abnormal Perception; *BIN1*-rs744373*G and *EPHA1*-rs11767557*C were associated with higher likelihood of Abnormal Perception; *NME8*-rs2718058*G was associated with a reduced likelihood Decreased Motivation. These findings suggest a common genetic etiology between MBI and traditionally recognized memory problems observed in AD and improve our understanding of the pathophysiological features underlying MBI.

## Introduction

Neuropsychiatric symptoms (NPS) are common to all types of neurocognitive disorders and are linked to poorer quality of life, more rapid functional decline, added disease burden, greater healthcare costs and higher rates of mortality^1^. There is increasing acknowledgment that NPS form an intrinsic aspect of prodromal dementia and may be an early marker of cognitive decline that can precede or accompany the onset of cognitive symptoms and clinical diagnosis^2^. The concept of ‘Mild Behavioral Impairment’ (MBI) describes the emergence of later-life NPS as an at-risk state for cognitive decline and dementia and as a potential manifestation of prodromal dementia^3^.

How NPS mechanistically link to the development of Mild Cognitive Impairment (MCI) and Alzheimer’s disease (AD) is not fully understood. Five potential models have been proposed^4,5^: 1) etiologic pathway where NPS have a direct deleterious effect on the brain leading to the AD symptoms; 2) shared genetic/environmental risk factors are causally related to the development of both cognitive impairment and NPS; 3) observed associations between NPS and MCI/dementia are a result of reverse causality, with individuals experiencing cognitive decline potentially showing reactive NPS; 4) synergistic reactions between NPS and other biological factors further increase the risk of developing MCI/dementia and 5) the accumulation of Alzheimer’s neuropathology and neurodengeration causes the genesis of psychiatric symptoms. Of interest to these five pathways is the role genetic risk loci for late onset Alzheimer’s Disease (LOAD) may play in the development of NPS and MBI, either as a shared genetic risk for both LOAD and NPS or by acting via intermediate pathways by promoting the accumulation of Alzheimer's neuropathology and subsequent NPS.

LOAD has a large genetic component, with genetic variants accounting for 53% of the total phenotypic variance of LOAD^6^. The *Apolipoprotein (APOE) epsilon 4* (*ε4) allele confers the largest known genetic risk for LOAD, with a 2-3 and 10-12 times increased risk for heterozygotes and homozygotes respectively^7^. Beyond *APOE*, genome wide association studies (GWAS), have identified single nucleotide polymorphisms (SNPs) at a further 23 loci associated with LOAD^8–13^ (Supplementary Table 1). These loci are clustered in biological pathways that are play an important role in the development of cognitive impairment and dementia, and are involved in the accumulation of the neuropathological features (amyloid beta (Aβ) and neurofibrillary tangles) of LOAD. The neuropathological features of LOAD have also been associated with greater impairment over time in NPS, and depression specifically, in cognitively normal and demented subjects^14–16^.

As such, LOAD risk loci may also be associated with the development of NPS, though, at present the role of LOAD risk loci in the development of NPS is unclear. Most research to date has focused on the association of *APOE* with individual NPS, with associations observed for depression, anxiety, apathy, delusions/hallucinations and agitation/aggression^17^. However, contrasting findings have been observed between studies on the association between *APOE* and affective symptoms (depression and anxiety), while there is limited cross-sectional evidence to suggest an association between apathy, agitation/aggression and delusions/hallucinations that has not been replicated in longitudinal studies^17^. To our knowledge, no study has examined the role of non-*APOE* LOAD risk loci with the development of NPS.

Here we report the associations of 24 genome-wide significant LOAD risk loci, either individually or collectively as a genetic risk score, with MBI in 1,226 sub-clinical and nondemented community dwelling adults.

## Methods

### Participants

Participants of this study were community dwelling older adults who were recruited into a longitudinal study of health and wellbeing, the Personality and Total Health Through Life project (PATH). The background and test procedures for PATH have been described in detail elsewhere^18^. We used data from the PATH Wave 4 60+ cohort. Briefly, at baseline (2001-2002) 2,551 older adults (60-64) were randomly sampled from the electoral rolls of Canberra and the neighboring town of Queanbeyan. 1,644 participants completed the fourth wave of assessments (2014-2015), of whom 1417 completed the informant based Neuropsychiatric Inventory. Individuals were excluded from analysis based on the following criteria: No available genomic DNA (n = 79); APOE ε2/ε4 genotype (n = 44, to avoid conflation of the ε2 protective and ε4 risk affect); non-European ancestry (n = 53); clinical diagnosis of dementia (n = 40); did not complete the Neuropsychiatric Inventory (n = 228); missing values in “Education” (total number of years of education assessed at wave 2, n = 18). This left a final sample of 1,226 individuals.

Written informed consent for participation in the PATH project was obtained from all participants according to the ‘National Statement’ guidelines of the National Health and Medical Research Council of Australia and following a protocol approved by the Human Research Ethics Committee of the Australian National University.

### Cognitive Function and Clinical Diagnosis

The diagnostic procedure used to determine cognitive function states and clinical diagnoses at Wave 4 are published elsewhere^2,19^. In summary, for the 1644 participants assessed at Wave 4, their data was screened for signs of decline based on the following criteria: a previous PATH diagnosis of dementia or a mild cognitive disorder, or evidence of current objective cognitive impairment (based on performance ≤ 6.7th percentile on at least one cognitive measure, or MMSE<24), and evidence of subjective decline on the Memory and Cognition Questionnaire MAC-Q^20^ or decline on the MMSE of >3 points since wave 3, or consistent MMSE<24 at waves 3 and 4^19^.

For participants who showed signs of cognitive impairment, as detailed above, an algorithm that combined neurocognitive assessments, informant data, and self-reported medical history was used to operationalize criteria for DSM-5 major NCD, mild NCD, DSM-IV dementia and MCI. Diagnoses were confirmed by a case file review by a research neurologist and consensus diagnosis with a clinician specializing in Psychiatry. Case files were reviewed by the research neurologist and included neuropsychological test data, informant data, structural brain MRI scans (where available) to aid differential diagnosis of dementia subtypes, selfreported medication list, and contact of participant for further clarification of details relevant to diagnosis^19^. Inter-rater reliability indicated high agreement between the neurologist and psychiatrist in the independent review of a subsample of 29 cases^19^.

MCI was assessed using the Winblad^21^ criteria for MCI. Subjects were considered ‘cognitively normal, but-at-risk’ (CN-AR) if they did not meet the Winblad criteria for MCI or the DSM-IV criteria for dementia; but demonstrated signs of impairment as identified by the screening criteria outlined above^19^. Individuals who did not meet any of the above criteria were classified as cognitively normal.

### Informant Interview

Informants were nominated by the PATH participant and provided information on the participant’s physical and mental health via a telephone interview. Informants were predominantly spouses (49.4%), children (33.8%), or a close friend (9.7%)^2^. The informant interview included the Neuropsychiatric Inventory (NPI)^22^, to assesses the presence and severity of dementia related behavioural symptoms, over one month, in 10 neuropsychiatric domains (delusions, hallucinations, agitation/aggression, dysphoria, anxiety, euphoria, apathy, disinhibition, irritability/lability, and aberrant motor activity) and two neurodegenerative domains (night-time behavioural disturbances and appetite and eating abnormalities).

### Mild Behavioral Impairment

The NPS Professional Interest Area (PIA) of the International Society to Advance Research and Treatment (ISTAART), a subgroup of the Alzheimer’s Association (AA), have developed and published research diagnostic criteria for MBI. According to the operationalized criteria^3^, MBI is hallmarked by changes in behavior or personality which start in later-life (after the age of 50 years), are sustained for at least six months, and represent a clear change from the person’s usual behavior or personality. The NPS of MBI have been clustered into the following domains: decreased motivation, affective dysregulation, impulse dyscontrol, social inappropriateness and abnormal perception or thought content. Importantly, these domains reflect areas of NPS shown to be valid and related to the syndromes of cognitive decline.

MBI was assessed in accordance with Criterion 1 of the ISTAART-AA diagnostic criteria for MBI^23^, but with a reference range of one month, using the Neuropsychiatric Inventory (NPI)^22^. A previously developed transformation matrix of the ISTAART-AA MBI criteria and the NPI domains was used to approximate whether individuals met the domain criteria for MBI^2,24^, as follows: 1) Decreased Motivation (NPI: apathy); 2) Affective/Emotional Dysregulation (NPI: dysphoria, anxiety, euphoria); 3) Impulse Dyscontrol (NPI: agitation/aggression, irritability/lability, aberrant motor behaviour); 4) social inappropriateness (NPI: disinhibition) and; 5) Abnormal Perception or Thought Content (NPI: delusions, hallucinations). The NPI neurovegetative domains are not reflected in the ISTAART-AA MBI criteria. Presence of at least one NPI behaviour symptom within a specific domain meant that the domain criteria was met.

### Genotyping and Genetic Risk Score

Genome-wide significant LOAD risk SNPs from 23 loci^8–13^ (*ABCA7, BIN1, CD2AP, CD33, CLU, CR1, EPHA1, MS4A4A, MS4A4E, MS4A6A, PICALM, HLA-DRB5, PTK2B, SORL1, SLC24A4-RIN3, DSG2, INPP5D, MEF2C, NME8, ZCWPW1, CELF1, FERMT2* and *CASS4;* Supplementary Table 1) were genotyped using TaqMan OpenArray assays as previously described^25,26^, in addition to the two SNPs defining the *APOE* alleles which were genotyped using TaqMan assays as previously described^27^. All SNPs were in Hardy-Weinberg equilibrium and genotype frequencies are reported in Supplementary Table 2.

For the genetic risk score analysis SNPs were coded additively according to the number of risk alleles, whereas for the single SNP analysis variants were coded additively according to the number of minor alleles (Supplementary Table 1). The *APOE* *ε2 and *APOE* *ε*4* alleles were assumed to be dominant to the *APOE* *ε3 allele. *APOE* alleles were coded as *APOE* *ε2+ (*APOE* *ε2/ε3 + *APOE* *ε2/ε2), *APOE* *ε*4*+ (*APOE* *ε*4*/ε*3* + *APOE* *ε*4*/ε*4*) or *APOE* *ε3/ε3. Participants with the *APOE* *ε2/ε4 allele were excluded to avoid conflation between the *APOE* *ε2 protective and *APOE* *ε4 risk effects.

Using the LOAD risk SNPs, an OR weighted genetic risk score (OR-GRS) was constructed, which is the sum of all the risk alleles across the individual, weighted by the Odds Ratio^28^. The OR-GRS is calculated according to the following formula: 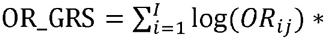 *G_ij_* for the *i*th patient, where log(*OR_ij_*)= the log of the odds ratio for the *j*th SNP and *G_ij_* = the number of risk alleles for *j*th SNP. GRS were not calculated for individuals with missing genetic data (n = 93). We weighted the LOAD risk SNPs using the previously reported OR for LOAD (Supplementary Table 1). As *APOE* is known to have the largest effect size for LOAD, the OR-GRS was also calculated excluding *APOE* to determine the effect of the GRS beyond that of *APOE*. The OR-GRS was transformed into a z-score.

### Data analysis

All analysis was performed in the R 3.3.2 Statistical computing environment. The association of the LOAD GRS with MBI domains was examined using logistic regression models adjusting for age, gender and years of education. Individual SNPs were similarly assessed, with each model only including a single SNP. Because 25 loci (*APOE* ε2, *APOE* ε4, & 23 LOAD GWAS SNPs) and one GRS were tested, p < 0.0019 was considered to be study wide significant after Bonferroni correction. p < 0.05 and >0. 0019 was considered nominally significant.

As a sensitivity analysis, the above models were re-run 1) with cognitive status included as a covariate and 2) with CN-AR and MCI participants excluded.

## Results

### Population characteristics of the PATH cohort

Descriptive characteristics of the wave 4 PATH cohort are presented in Table 1. The distribution of the OR-GRS is multimodal, with modes at 3.15, 4.60 and 5.58 corresponding to the presence of zero, one or two *APOE* ε4 alleles. The most prevalent MBI domains within the PATH cohort are Affective Dysregulation and Impulse Dyscontrol, affecting ~21% of the sample.

**Table 1:**
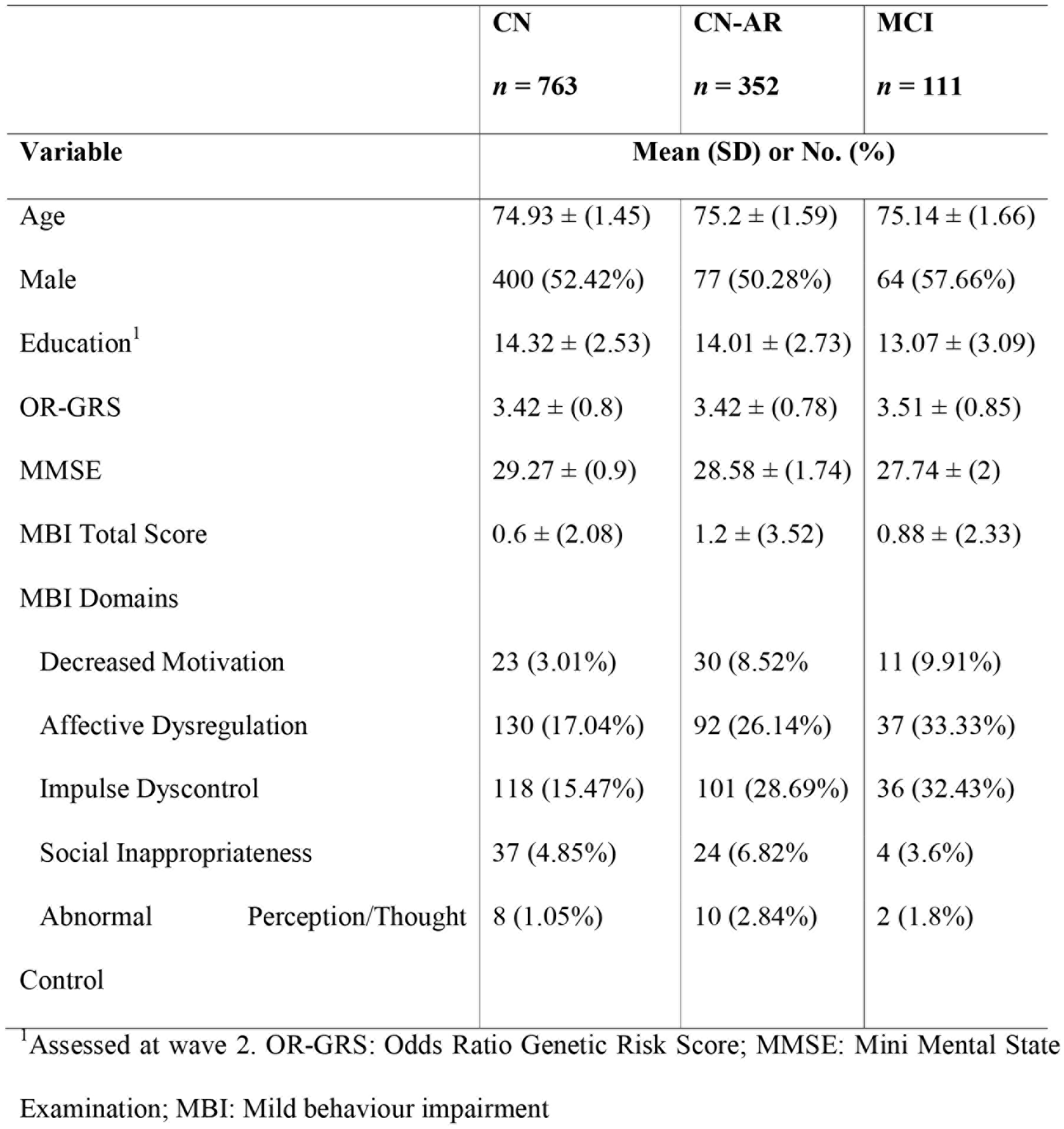
Descriptive Statistics of PATH cohort at Wave 4

### Association of AD related genetic variants with MBI

A binomial logistic regression was performed to evaluate the effects of the OR-GRS on the likelihood of participants exhibiting MBI symptoms (Table 2). Increasing OR-GRS was associated with an increased likelihood of exhibiting Affective Dysregulation, with a 1SD increase having a 1.23 higher odds (*p* = 0.0047). However, when *APOE* was excluded from the OR-GRS, it was no longer significantly associated with any of the MBI domains. We further evaluated the effects of the individual SNPs on MBI, the observed associations would not withstand correction for multiple testing and we report the results that were nominally significant (Table 2). The *APOE*\*ε4 allele was associated with an increased likelihood of Affective Dysregulation; *MS4A4A*\*C and *MS4A6A*\*G were associated a reduced likelihood of Affective Dysregulation; NME8*G was associated with a reduced likelihood Decreased Motivation; *ZCWPW1*\*C was associated with a reduced likelihood Social Inappropriateness and Abnormal Perception/Thoughts; *BIN1*\*G and *EPHA1*\*C were associated with an increased likelihood of Abnormal Perception/Thoughts. The effect of the OR-GRS or individual SNPs on the likelihood of exhibiting behavioural symptoms as assessed by the NPI are presented in Supplementary Table 3. These results are largely non-significant and would not withstand correction for multiple testing.

**Table 2:**
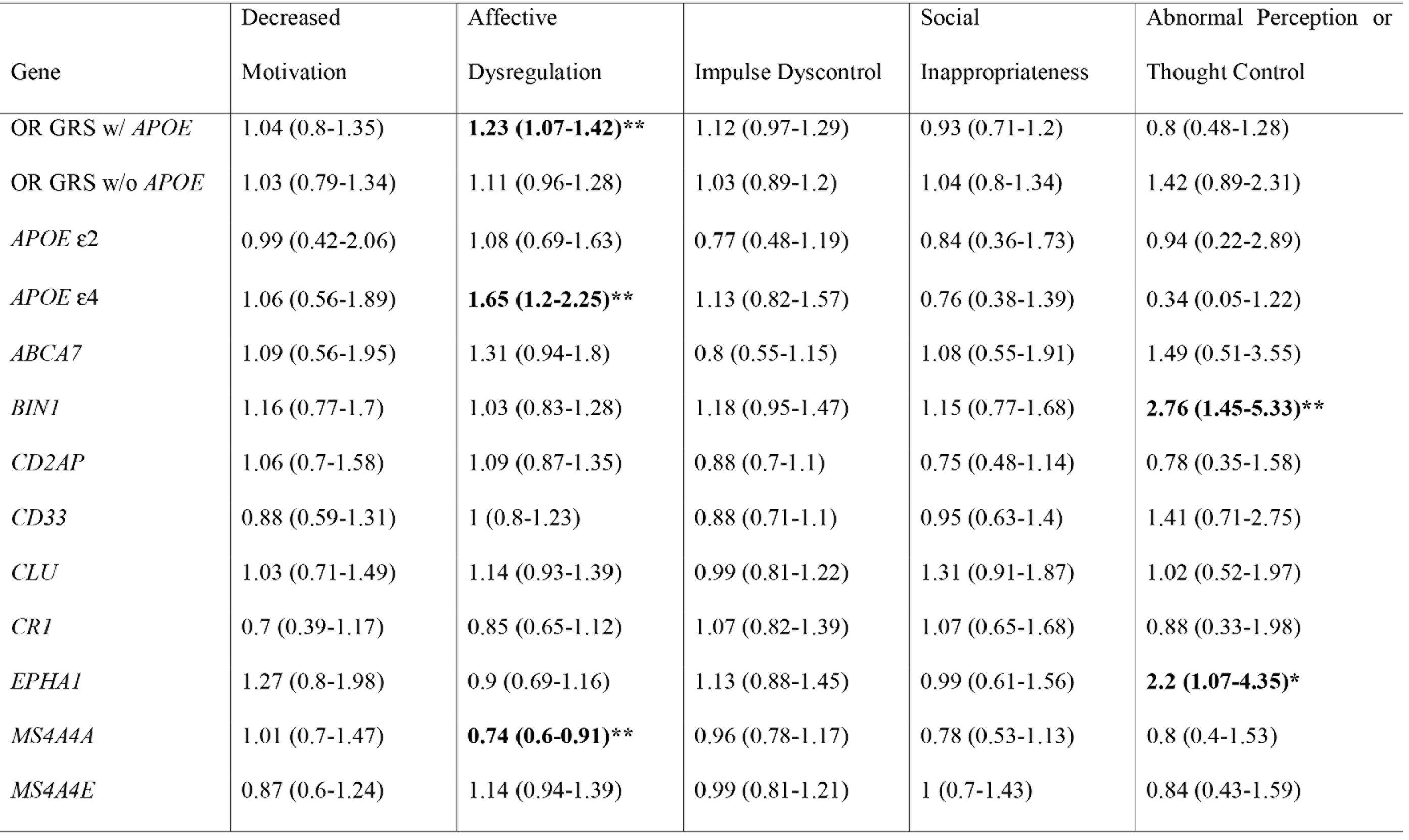
Association of AD genetic risk loci with Mild Behavioural Impairment domains

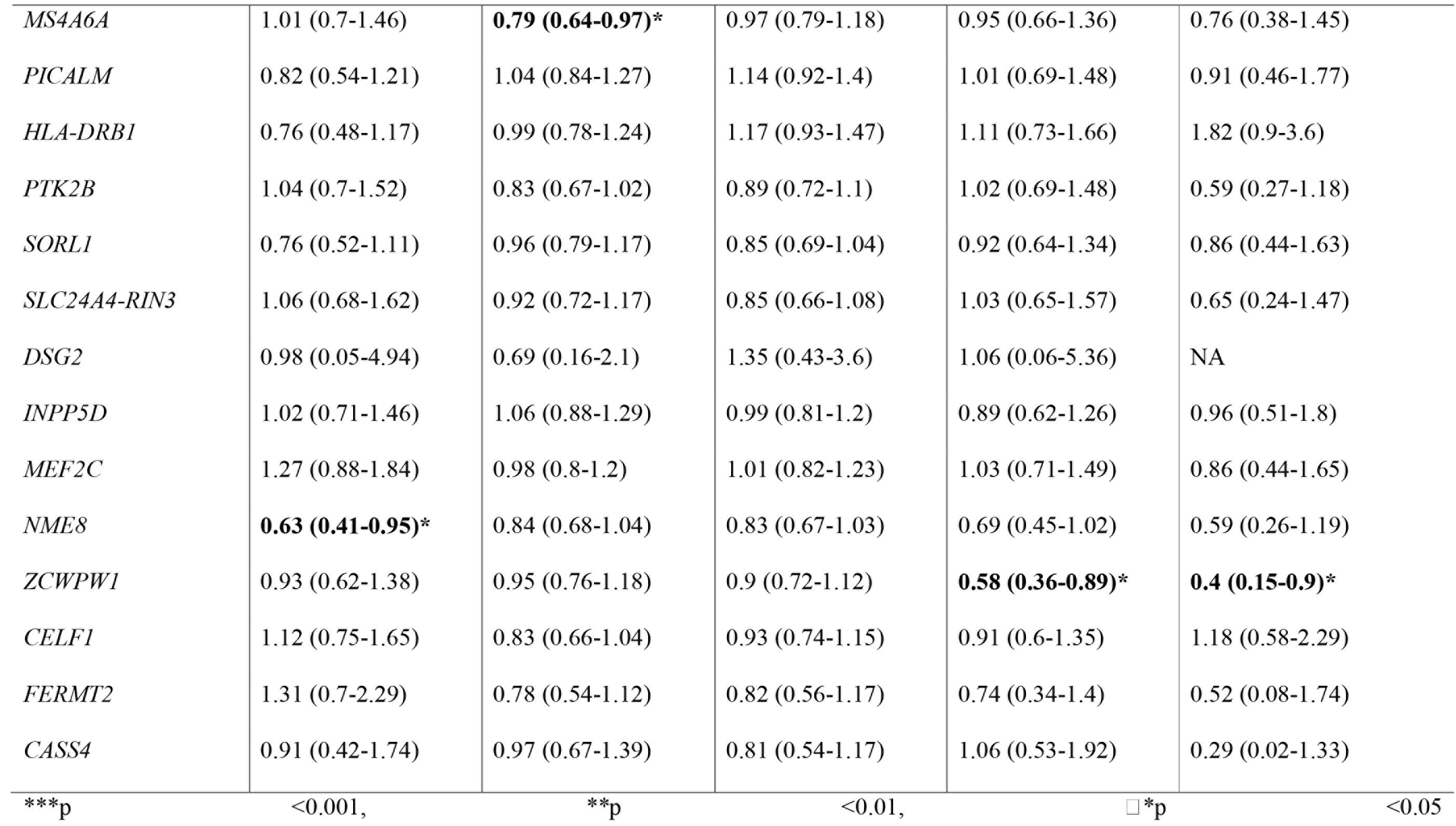

Including cognitive status as a covariate in the models did not markedly change the results (Supplementary Table 4) for the association of the AD genetic loci with MBI. When assessing the associations in CN participants only (Supplementary Table 5), the results were largely similar, though some differences were observed, with ZCWPW1*C no longer associated with Abnormal Perception/Thought Control; *FERMT2*\*C been associated with a decreased risk of impulse dyscontrol and; *HLA-DRB1*\*T associated with an increased risk of impulse dyscontrol and Abnormal Perception/Thoughts.

## Discussion

In this study, we investigated the association of genome-wide significant LOAD risk loci with MBI symptoms in a large cohort of cognitively normal and sub-clinical participants. We identified 3 loci (*APOE, MS4A4A, MS4A6A*) associated with Affective Dysregulation, 1 loci (*NME8*) associated with Decreased Motivation, 1 loci (ZCWPW1) associated with Social Inappropriateness and three loci associated with (*BIN1, EPHA1, ZCWPW1*) Abnormal Perception/Thought Control. In addition, a weighted GRS was associated with Affective Dysregulation. However, this association was attenuated when *APOE* was excluded from the GRS, indicating that the association was driven by the dominant effect of the *APOE*\*ε4 allele. The majority of these findings, however, were nominally significant (p < 0.05 and >0.0019), with only the *APOE* - affective dysregulation association significant after multiple testing (*p* = 0.0018).

The association between *APOE* and depression in population based studies and in dementia patients has been widely investigated in the literature, however, the results between studies are mixed^17,29^. A recent meta-analysis including 13 studies of late-life depression found that the APOE ε4 allele significantly increased risk of depression^30^. Furthermore, a longitudinal population based analysis found that the *APOE* ε4 allele was associated with incident minor depression and depression symptom severity over 5 years, even after excluding participants who developed dementia within 9 years^29^. The results from our analysis provide additional support for *APOE* promoting increased risk of late-life depression, though further longitudinal analysis is needed to verify these findings. This additional analysis is needed as it remains unclear whether depression is a risk factor of AD and dementia or an early manifestation of AD brain pathology^1^.

The mechanisms underlying the increased risk of depression in *APOE* ε4 carriers are not fully understood, with several potential mechanisms been implicated. First, brain atrophy may act as an intermediating factor, with *APOE* e4 carriers exhibiting greater medial temporal lobe atrophy^31^. Temporal lobe atrophy has also been associated an increased risk of incident major depression independently of dementia^32^. Second, *APOE* is associated with cerebrovascular dysfunction^33^, while late-life depression is associated with increased cerebrovascular comorbidities and microvascular lesions^34^. This suggests that cerebrovascular dysfunction induced by the *APOE* ε4 may increase cerebrovascular damage leading to increased depressive symptoms. Third, *APOE* influences amyloid-β (Aβ) aggregation, deposition and clearance, with the ε4 allele associated with increased Aβ levels and plaque burden^31^. Increased Aβ levels have also been associated with depression and worsening depressive symptoms over time^14,35^. This maybe driven by a neuroinflammatory response as a result of microglial activation by Aβ^36^, promoting the release of inflammatory cytokines that interferes with neurotransmitters and neurocircuitry, leading to depressive symptoms^37^.

In addition to *APOE*, two SNPs within the *MS4A* locus were associated with a decreased risk of affective dysregulation. The function of the proteins encoded by the genes within the *MS4A* gene cluster is not well characterized, however recent reports have suggested a putative role in the immune system by promoting activation of microglia and the release of pro-inflammation of cytokines^38–40^. Inflammation impacts the synaptic availability of the monoamines serotonin, noradrenaline and dopamine, as well as the excitatory amino acid glutamate, that can ultimately affect the neurocircuitry that regulates behavior associated with anhedonia and anxiety, core aspects of depression^37^. As such, the *MS4A* putative role in the immune system may influence both AD pathology and depressive symptoms.

*BIN1* and *EPHA1* were both associated with the MBI domain abnormal perception/thought control. In AD patients with psychosis (AD + P), neuroimaging and post-mortem data have indicated an exaggerated prefrontal cortical synaptic deficit^41^. Greater synapse loss in AD + P maybe a result of either increased accumulation of pathology in AD + P or from enhanced synaptic vulnerability to these pathology due to other molecular changes ^41^. In agreement with this, tau pathology has been consistently shown to be increased in AD + P ^41,42^ The association between *BIN1* and tau pathology has been firmly established, where *BIN1* knockdown in a Drosophila model suppressed tau-mediated neurotoxicity, while in human neuroblastoma cell lines and the mouse brain BIN1 and tau were observed to colocalize and interact^43^. Accordingly, *BIN1* has been associated increased neurofibrillary tangle burden^44^. The results from this analysis suggests that *BIN1* may influence tau pathological burden, promoting synaptic loss and the development of psychotic symptoms within the prodromal phase of dementia. The role of *EPHA1* in AD is not well understood, however, it is highly expressed in the adult brain and plays a role in synaptic formation and plasticity, axonal guidance and brain development^45–47^. This suggests that variants in *EPHA1* may enhance synaptic vulnerability to AD pathology, and corresponding psychotic symptoms. However, these results should be interpreted with caution. A recent genome wide association study of AD + P did find that both *BIN1* and *EPHA1* were associated with AD + P when contrasting to controls, however no association was observed when contrasting AD + P to AD - P^48^. This suggests that both loci are associated with AD irrespective of psychotic symptoms, though this analysis did have limited power.

Finally, we observed a decreased risk for NME8 with decreased motivation and ZCWPW1 with a decreased risk of social inappropriateness and abnormal perception/thought control. The role of *NME8* and *ZCWPW1* in AD is not well characterised, with both loci relatively understudied^49^. *NME8* has been previously associated with non-neurological related diseases^50^, and recently with cognitive decline, elevated CSF tau and hippocampal atrophy^50^. The pathophysiology of apathy in AD is characterised by dysfunctions in the prefrontal cortex, orbitofrontal cortex, anterior cingulate cortex, amygdala and basal ganglia, particularly in regard to cortico-subcortical circuits involving dopaminergic and cholinergic pathways^51,52^. As with *NME8*, the function of *ZCWPW1* is unknown, though the index SNP was shown to have functional evidence as an expression quantitative trait locus for *PILRB*, which is expressed in microglia and is involved in the regulation of immune response^53^. The neurobiological correlates of disinhibition in AD include Orbitofrontal-subcortical circuit dysfunction, that impairs social cognitive abilities and loss of control over reactions^52,54^. Due to the scarcity of studies investigating the role of *NME8* and ZCWPW1, the possible underlying mechanisms for their associations with the MBI domains are not known.

The results from this study should be interpreted in conjunctions with some study limitations. First, as this is a candidate gene study, which can be subject to false positive findings, further replication of our results is needed in an independent cohort. Second, MBI was assessed using the NPI rather than the recently published MBI checklist. The NPI rating scales are designed to assess neuropsychiatric symptoms in dementia, and as such might not be sensitive to milder symptomology in functionally independent community dwelling adults^23^. Additionally, the NPI assesses neuropsychiatric symptoms over a short reference time periods that maybe confounded by transient reactive states (e.g. sleep deprivation, medications, adversity) when used in the context of prodromal states^23^. Third, as the NPS are assessed via informants, the neuropsychiatric data maybe susceptible to recall bias, influenced by the informants mood, cultural beliefs, denial or education^55^. Finally, as the neuropsychiatric data were only collected at wave 4 in PATH, we are unable to conduct longitudinal analysis to assess whether the AD risk loci are associated with progression in NPS. Despite these limitations, this study had several strengths, including consisting of a large population based sample, inclusion of all the known GWAS LOAD risk loci and the narrow age-range reduces the influence of age differences on the results.

In conclusion, this is the first study to investigate the association of LOAD genetic risk loci with MBI. We found that five LOAD risk loci (*APOE, MS4A, BIN1, EPHA1, NME8* and *ZCWPW1*) are associated with MBI domains. Nevertheless, the results from this study need to be replicated in independent cohorts to validate our findings, as the *APOE* - affective dysregulation association is the only test to survive correction for multiple testing. These findings suggest a common genetic etiology between MBI and traditionally recognized memory problems observed in dementia/AD and improve our understanding of the pathophysiological features underlying MBI.

## Acknowledgments

We thank the investigators in the PATH study: Nicolas Cherbuin, Peter Butterworth, Andrew Mackinnon, Anthony Jorm, Bryan Rodgers, Helen Christensen, Patricia Jacomb, Karen Maxwell and Simon Easteal. We also thank the PATH participants. The study was supported by the National Health and Medical Research Council (NHMRC) grants 179805 and 1002160 and the NHMRC Dementia Collaborative Research Centre Early Diagnosis and Prevention. SJA is funded by the ARC Centre of Excellence in Population Ageing Research, ARC grant CE1101029. KJA is funded by NHMRC Research Fellowship No. 1002560. ZI is funded by the Hotchkiss Brain Institute with support from the Alzheimer Society Calgary. MEM is supported by the Australian National Health and Medical Research Council (NHMRC) and Australian Research Council (ARC) Dementia Research Development Fellowship #1102028.

